# *Burkholderia pseudomallei* rubrerythrin promiscuously binds metals in a structurally pre-formed bimetallic binding site

**DOI:** 10.1101/2025.06.01.657255

**Authors:** Gabrielle R Budziszewski, Miranda L Lynch, M Elizabeth Snell, Diana CF Monteiro, Sarah EJ Bowman

## Abstract

Rubrerythrins are a group of proteins within the Ferritin-like superfamily that display a defining four-helix bundle domain. They also show multiple structural features that are crucial to their functionality as iron storage proteins and in detoxification and oxidative stress response. Here we investigate rubrerythrin (Rbr) in multiple metalated states, from the pathogen *Burkholderia pseudomallei* (*Bp*). We use X-ray crystallography for structure determination of Rbr to probe the capacity and specificity of metal binding. *Bp*Rbr lacks the rubredoxin moiety found in canonical Rbrs from anaerobic lineages, and we demonstrate that *Bp*Rbr also possesses a domain-swapped dimer, which has functional implications for its putative role in oxidative stress response. We also carry out *in crystallo* spectroscopic assessment of *Bp*Rbr with various metals, using energy dispersive X-ray (EDX) spectroscopy. We observe that samples can contain metals other than those supplied in crystallization conditions, and developed a strategy of utilizing EDX spectroscopy to select those samples with single metal incorporation for downstream diffraction data collection. Our work underscores the importance of spectroscopic probing for definitive metal identification and characterization.

**Highlights:**

- Present structures of apo-, Fe-, Mn-, and Co-bound rubrerythrin from *B. pseudomallei*
- *Bp*Rbr lacks rubredoxin domain and possesses unique domain swap fold in the dimer
- *Bp*Rbr features include promiscuous metal binding and pre-formed metal binding sites
- Highlights importance of *in crystallo* spectroscopy in investigating metalloproteins
- Mutant Rbr with abrogated metal binding supports premise of pre-formed binding sites

## Introduction

Increasing levels of antibacterial resistance in microorganisms coupled with amplified risks associated with emerging pathogens present growing challenges to human health (Casadevall, 2020, Dolecek *et al*., 2022, Mora *et al*., 2022). This work focuses on a protein from the pathogen *Burkholderia pseudomallei* (*Bp*), a gram negative, aerobic soil bacterium that is intrinsically resistant to many antibiotics. Indeed, the entire *Burkholderia* genus contains multiple species that are opportunistic pathogens of both humans and other mammals. Some *Burkholderia* species are especially serious for immunocompromised patients, most notably those with cystic fibrosis, diabetes, and chronic kidney disease (Casadevall, 2020). *Bp* is the causative agent of melioidosis, a life-threatening illness that can impact organ function and induce septic shock, and is spread via inhalation, ingestion, or contact through broken skin with contaminated soil or water (Phillips & Garcia, 2024). The mortality rate has been estimated as ∼10-40% of infected melioidosis patients, depending on correct diagnosis, comorbidities, and other risk factors (Phillips & Garcia, 2024). These factors make *B. pseudomallei* a dangerous emerging threat; it is classified as a Tier 1 Select Agent by the Centers for Disease Control and is considered a potential bioterror agent due to the high potential for aerosol spread and its antimicrobial resistance (Schweizer, 2012). Worryingly, cases of melioidosis have been reported recently in the Gulf Coast of Mississippi and other non-endemic regions, highlighting the need for further study of *B. pseudomallei* as an emerging pathogen (Petras *et al*., 2023, Torres, 2023).

One strategy proposed to target pathogenic bacteria such as *B. pseudomallei* is to focus on proteins that function in oxidative stress response (OSR) pathways, which allow bacteria to adjust to different environmental circumstances, such as changes in oxygen levels, chemical conditions, salinity, and pH, among other features (Vatansever *et al*., 2013, Dawan & Ahn, 2022). These pathways enable a robust response to withstand the onslaught of the human innate immune system, which uses reactive oxygen species such as hydrogen peroxide to attack pathogens. Furthermore, existing antibiotic treatments can induce oxidative stress, making the antioxidant system a key target to sensitize bacteria to known treatments. One such antioxidant protein in *B. pseudomallei* is rubrerythrin (*Bp*Rbr), a metal-binding protein with putative function in oxidative stress response. *Bp* has a relatively large genome (7.24 Mb) organized on two chromosomes (4.07Mb and 3.17Mb, respectively), with extensive genomic plasticity, which potentially contributes to its ability to evade immune response (Holden *et al*., 2004). Many of the *Bp* genes associated with proteins expressed in response to environmental stress are primarily located on chromosome two. The operon that encodes *Bp*Rbr is on chromosome two; rubrerythrin (Rbr) has been shown to be upregulated as part of the OSR in *B. pseudomallei* and in other pathogens (Sztukowska *et al*., 2002, Jitprasutwit *et al*., 2014). Interestingly, it has also been shown that metal acquisition plays a role in *Bp* virulence and pathogenicity (Butt & Thomas, 2017).

Here we investigate Rbr from the aerobic bacterium *B. pseudomallei*. In anaerobic bacteria, Rbr proteins have been shown to have peroxidase activity with a di-iron metal active site and a rubredoxin domain (Coulter *et al*., 1999, Weinberg *et al*., 2004, Barreiro *et al*., 2023, Dillard *et al*., 2011). In the anaerobes *Pyroccocus furiosus* and *Nitratidesulfovibrio vulgaris* (formerly *Desulfovibrio vulgaris*), the Rbr operon is predicted to contain additional metal-dependent proteins (Fig 1). A homologous protein, sulerythrin (SulE) from the aerobe *Sulfolobus tokodaii* exhibits catalase and peroxidase activity (Jeoung *et al*., 2021). Rbr proteins are in the ferritin-like superfamily, with a four-helix bundle fold around the di-iron substrate binding site. Canonical Rbr proteins found in anaerobic lineages are functionally dimeric, with each monomer having a Rubredoxin domain that is involved in its peroxidase activity. Previous structural and phylogenetic studies have identified that *Bp*Rbr differs from Rbrs from anaerobic lineages. It was determined that Rbr proteins in most aerobic bacteria do not contain the rubredoxin domain (Cardenas *et al*., 2016). One previous structure of *Bp*Rbr has been deposited to the Protein Data Bank (PDB 4DI0) by the Seattle Structural Genomics Center for Infectious Disease (SSGCID, 2012), but no additional functional or structural studies of this protein from *Bp* have been reported. Unlike many Rbr proteins, which form dimers in which a single chain contributes all four helices to the four-helix bundle (Cardenas *et al*., 2016), *Bp*Rbr exhibits domain swapping, with each chain contributing two helices to each four-helix bundle. We are especially interested in the structure and function of *Bp*Rbr from this aerobic species, as we hypothesize that the exhibited domain swapping may be correlated with evolutionary exposure to oxygen. *Bp*Rbr is annotated as a metal binding protein that may function as an oxidoreductase. The protein has been identified as upregulated in response to oxidative stress (Jitprasutwit *et al*., 2014) and impacting virulence (Burkholderia Genome DB, 2024).

**Figure 1:**
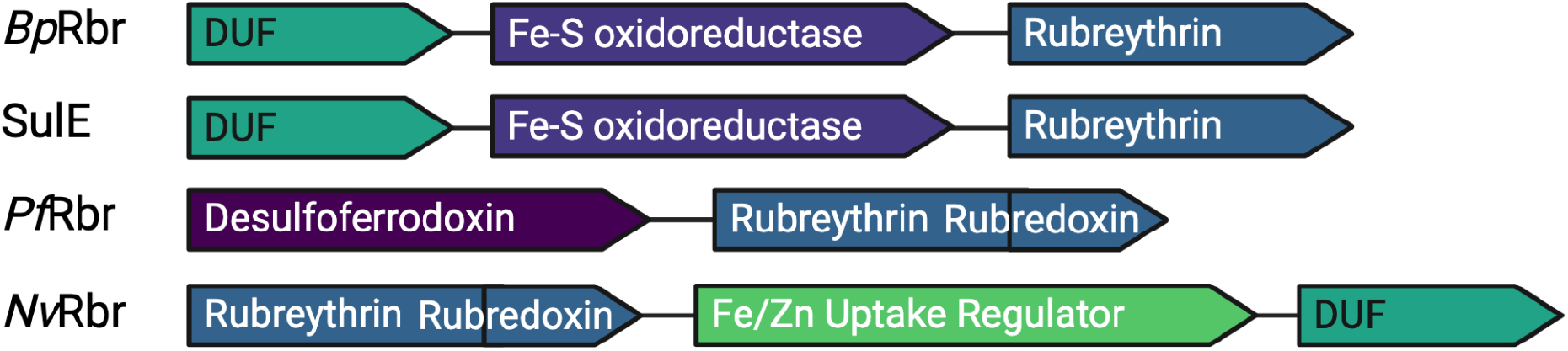
Operon organization for rubrerythrin family of proteins from different bacterial species. DUF represents a domain of unknown function.

In this work, we present a collection of structures of multiple *Bp*Rbr constructs in apo- and metal-bound forms, including a mutant form that does not bind metal in the di-metal site. The apo-structure is not fully demetalated, and so we refer to this as quasi-apo (*Bp*Rbr_q-apo_). The structures have been obtained with a combined approach that uses *in crystallo* EDX spectroscopy as a selection tool for correctly metalated crystals, and then X-ray crystallography for diffraction data collection for structure determination. EDX spectroscopy measurements upstream of data collection were carried out to ensure samples had metal-specific single metal incorporation into the di-metal site, as our investigations have revealed that multiple metals can bind even if they are not supplied in the crystallization conditions. The structures are similar across the series from *Bp*Rbr_q-apo_ to metal bound wild-type and mutant, suggesting that the metal binding site is pre-formed. Our studies demonstrate the capabilities of *Bp*Rbr for promiscuous metal binding, the presence of the organized binding site even in the absence of metal binding, and the importance of spectroscopic probing of metal-containing samples prior to X-ray diffraction for definitive metal identification and characterization.

## Materials and Methods

### Operon mapping for BpRbr

The sequence from *Burkholderia pseudomallei* 1710b chromosome II genome was obtained from GenBank (Woods & Nierman, 2005); sequences from *S. tokodaii, P. furiosus*, and *N. vulgaris* were obtained from NCBI. Operon organization for each genomic sequence was obtained using the Web-based program Operon Mapper (Taboada *et al*., 2018). Both *B. pseudomallei* and *S. tokodaii*, the two aerobic species, show high probability that the Rbr gene is on an operon with a domain of unknown function (DUF) hypothetical protein in Rubrerythrin cluster and an Fe-S oxidoreductase-like protein. In the anaerobic bacteria, other genes predicted to be on the Rbr operon include a desulfoferrodoxin (in *P. furiosus*) and a Fur/Zur metal uptake regulator (in *N. vulgaris*).

### Construct design

Plasmid with *Bp*Rbr gene was obtained from VectorBuilder. The *Bp*Rbr gene from *B. pseudomallei* (uniprot code Q3JK21), bearing an N-terminal tail with a TEV cleavage site (ENLYFQ↑GS) and BamHI cleavage sequence was cloned into a PET28a plasmid using Ndel and XhoI cloning sites (pET28a-Rbr plasmid). The expression product contained the N-terminal tail MGSSHHHHHHSSGLVPRGSHMENLYFQGS. The plasmid was propagated by transformation into *E. coli* XL1 blue cells (Agilent). Generation of the non-metal binding mutant was rationally designed by mutation of glutamate residues that bridge both metal sites, E53 and E124. The *Bp*Rbr-E53A/E124A mutant (*Bp*Rbr_mut_) was generated by site-directed mutagenesis and its sequence was verified by Sanger sequencing.

### Protein expression and purification

Hexahistidine-tagged *Bp*Rbr was expressed from commercial *E. coli* BL21 DE3 Gold cells (Agilent). The cells were transformed with the pET28a-His-*Bp*Rbr plasmid and a starter culture (20 mL) in LB medium was grown overnight at 37 ºC with agitation (200 rpm) with kanamycin selection. 1L auto-induction media with kanamycin (50 ug/L) was inoculated with 10 mL of overnight culture, grown at 37 ºC for 4 hours followed by 16 h at 22 ºC with agitation (200 rpm). Cells were collected by centrifugation (10 kg, 15 min), suspended in lysis buffer (50 mM Tris pH 7.5, 300 mM NaCl, 10 mM imidazole, 1 mM TCEP) disrupted by sonication and the cell debris separated by centrifugation (30 kg, 45 min). *Bp*Rbr was bound to Ni-NTA resin, the resin washed with 50 mL of wash buffer (lysis buffer with 50 mM imidazole) and *Bp*Rbr eluted in 10 mL fractions with elution buffer (lysis buffer with 300 mM imidazole). Fractions containing *Bp*Rbr were identified by SDS-PAGE, and pooled. For tag removal, pooled material from Ni-NTA purification was dialyzed overnight into lysis buffer in the presence of 2% molar equivalents of TEV at 4 ºC. Efficacy of TEV cleavage was assessed by SDS PAGE and the protein was subjected to reverse Ni-NTA purification. Following reverse Ni-NTA purification, protein was dialyzed into 20 mM HEPES, pH 8.0, 100 mM NaCl, 1 mM TCEP, 10 mM EDTA with two buffer changes to chelate metals acquired during purification. Protein was subsequently loaded into a Superdex S75 16-600 SEC column (Cytiva) and isocratically eluted (20 mM HEPES pH 8.0, 100 mM NaCl, 1 mM TCEP). *Bp*Rbr containing fractions were collected, the protein concentrated and flash frozen for storage. Typical yields were of 50 mg/L of culture. *Bp*Rbr_mut_ was similarly expressed and purified. Tagged *Bp*Rbr was also similarly expressed and purified, except that the TEV cleavage and demetalation steps were skipped and the protein purified to homogeneity directly by gel filtration following the initial Ni-NTA purification.

### Protein crystallization

Initial crystallization hits were determined from high-throughput screening at the National Crystallization Center at UB HWI (Budziszewski *et al*., 2023, Lynch *et al*., 2023). *Bp*Rbr protein concentration for crystallization was 10-15 mg/mL. Optimization was performed by matrix screening in 96-well plates in 1 μL sitting drops using vapor diffusion for apo- and Mn-*Bp*Rbr and microbatch-under-oil using paraffin oil (EMD Chemicals) for Fe-*Bp*Rbr. Mn- and Fe-*Bp*Rbr crystals were generated from seeds from apo crystals. The final crystallization conditions for each sample were as follows: *Bp*Rbr_q-apo_ 0.15 M Li2SO4, 0.1 M Bis-tris propane, pH 7.0; Mn-*Bp*Rbr 0.14M Li_2_SO_4_, 0.09 M Bis-tris propane, pH 7.75, 22% PEG 3350, 0.8 mM MnCl_2_; Fe-*Bp*Rbr 0.15 M Li_2_SO_4_, 0.1 M Bis-tris propane, pH 6.75, 30% PEG 3350, 1 mM FeCl_2_. Apo- and Mn-*Bp*Rbr crystals were treated with PEG 400 as a cryoprotectant, then harvested and vitrified in liquid nitrogen. Fe-*Bp*Rbr was harvested through the oil as cryoprotectant and vitrified in liquid nitrogen. Apo crystals for metal soaks of untagged *Bp*Rbr were grown in vapor diffusion sitting drop format in 10 mM HEPES pH 7.5, 0.1 M lithium sulfate, 23% PEG 3350. Crystals were soaked in a solution of mother liquor supplemented with 1 mM metal of interest and 30% glycerol and vitrified in liquid nitrogen prior to data collection. *Bp*Rbr_mut_ was crystallized in a vapor diffusion sitting drop format at room temperature in 0.1 M sodium citrate, pH 5.0, 0.1 M potassium thiocyanate, 22% PEG 4000. *Bp*Rbr_mut_ crystal used for structure determination was cryoprotected with 1.5 M hexanediol.

### EDX data collection

Prior to X-ray diffraction data collection, EDX spectra were collected for each crystal sample at SSRL BL 9-2 using the Blu-Ice excitation scan mode (Soltis *et al*., 2008). The nickel absorption K-edge energy (8333 eV) was used for all samples, with a 10 s scan for each crystal. The EDX spectra produce K_α_ and K_β_ emission lines that enable metal identification, as each element has specific emission transitions that align with fluorescent counts (Thompson *et al*., 2009). The EDX data were plotted using R (R Core Team, 2024) and the viridis(Lite) palette (Garnier *et al*., 2024).

### X-ray data collection, structure determination and refinement

Diffraction data for *Bp*Rbr_q-apo_ were collected at NSLS-II at the FMX beamline 17-ID-2 (4 × 10^12^ ph/s, 0.97933 Å, 100K) using a Dectris Pilatus 6M detector. Reflections were indexed, integrated and scaled using autoproc (Vonrhein *et al*., 2011, Evans, 2006, Evans & Murshudov, 2013, Winn *et al*., 2011). Due to anisotropy, the data were truncated in an ellipsoidal fashion where local(I/sigI) >= 1.2 using StarAniso (Tickle *et al*., 2018). Data were phased using the molecular replacement program MOLREP (Vagin & Teplyakov, 1997) using chain A of our original *Bp*Rbr structure as a search model (PDB ID: 8FUH). Refinement was carried out in both refmac5 and phenix refine (Murshudov *et al*., 2011, Liebschner *et al*., 2019). Metal occupancies were allowed to refine in phenix refine and adjusted manually to reflect both 2F_o_-F_c_ and F_o_-F_c_ difference maps. Polder maps were generated using the phenix software suite (Liebschner *et al*., 2017).

Diffraction data for metal-soaked *Bp*Rbr crystals were collected at SSRL beamline 9-2 (6.4 × 10^11^ph/s, 0.97946 Å, 100K) using a Dectris Pilatus 6M detector. EDX scans were performed prior to X-ray data collection at SSRL beamline 9-2 or 12-1 by adjusting the energy to the nickel K-edge (8332.8 eV). Data from a 180 degree rotation were collected in 0.2 degree increments between frames at a detector distance of 250 mm. Reflections were indexed, integrated and scaled using autoXDS and aimless (Kabsch, 2010). Only datasets indexed into R3(H3) were pursued for structure solution to retain consistency through dataset comparisons. Data processing and structure solution were carried out as described for other *Bp*Rbr_q-apo_.

Diffraction data for the *Bp*Rbr_mut_ mutant structure were collected at NSLS-II at the AMX beamline 17-ID-1 (4.3 × 10^12^ ph/s, 0.920100 Å, 100 K) using an Eiger 9M detector. Data from a 180 degree rotation were collected in 0.2 degree increments between frames at a detector distance of 120 mm. Data processing and structure solution were carried out as described for other *Bp*Rbr structures.

## Results and Discussion

### Structures of BpRbr

We have solved several structures of *Bp*Rbr for q-apo- and metal-bound constructs. All structures were solved in space group R3, showing six copies of the protein in the asymmetric unit (Fig 2A), arranged as three domain swapped dimers. The dimer is clearly identifiable (Fig 2B, colored as dark blue and cyan). Size exclusion chromatographs obtained during the last purification step showed a single peak with a retention time matching a dimer, rather than a hexamer. Two putative metal-binding sites are observed in each four-helix bundle, which are generated by domain swapping of two helices from each monomer unit (Fig 2C). The di-metal sites of *Bp*Rbr consist of each metal site coordinated by a glutamate-glutamate-histidine motif. Two of the glutamate residues (E53 and E124) bridge the two sites; in site one the coordinating residues are E20, H56 and the two bridging glutamates and in site two the coordinating residues are E90, H127 and the two bridging glutamates (Fig 2C). One of the axial metal coordination sites is occupied by the histidine residue in each coordination center, and the second axial site is a coordinating water (Fig 2D). Two four-helix bundles comprise each dimer, and each four-helix bundle has two sites for metal binding (i.e. there are four metal binding sites per dimer). Above the putative substrate binding pocket, one or more water molecules are observed, capped by E93 (Fig 2D).

**Figure 2:**
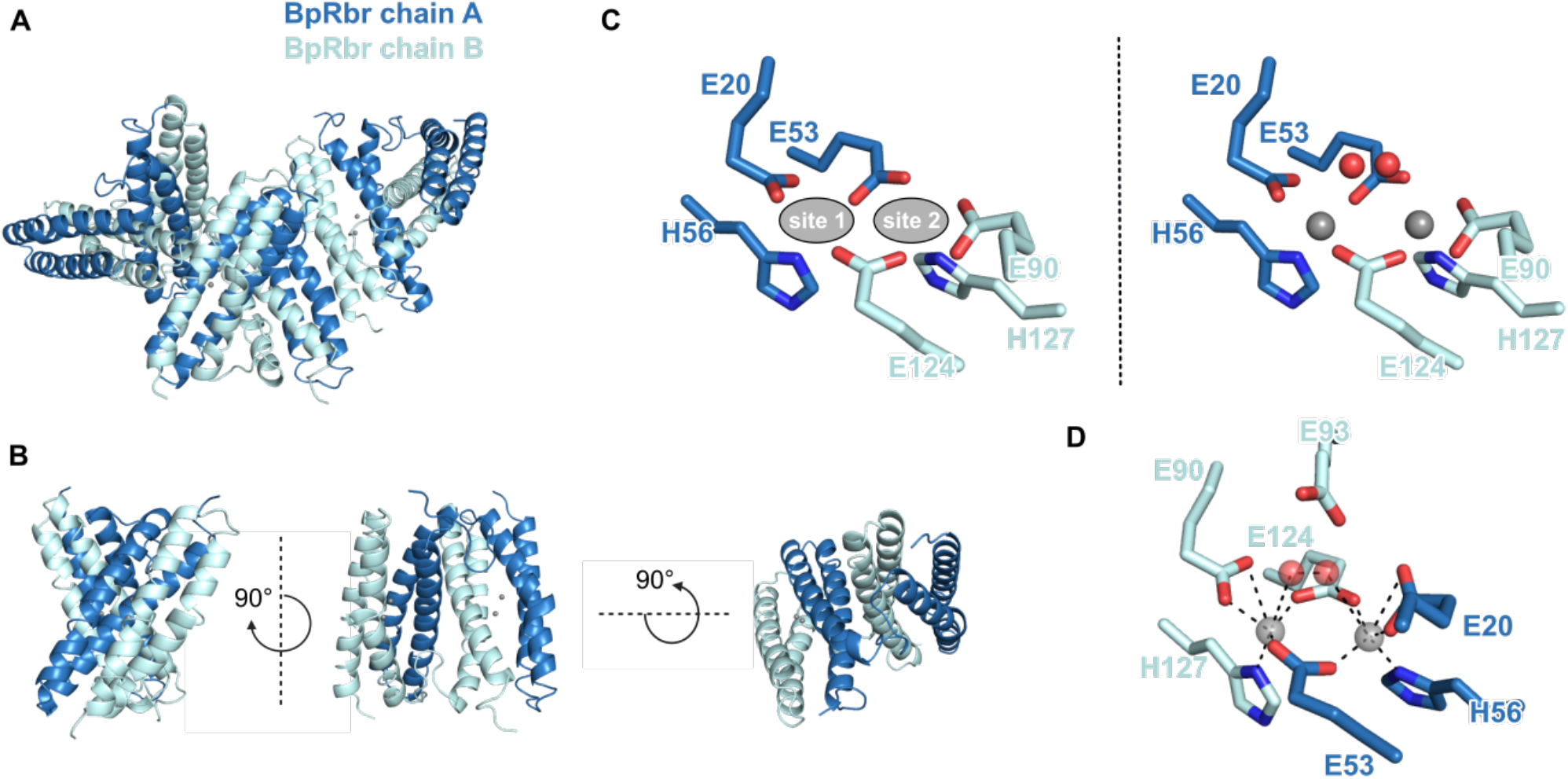
Overall structural model of *Bp*Rbr (PDB 9ONM). A) The asymmetric unit cell contains three *Bp*Rbr dimers. B) *Bp*Rbr dimer in a domain-swapped conformation. C) Left: Dual metal-binding sites formed at each four-helix bundle consist of a site 1 and site 2 composed of an EEH motif. Right: Site 1 and site 2 metal positions are represented by grey spheres and a dynamic solvent molecule is represented by two red spheres. D) Mobile solvent molecules above the binding site provide another coordination site for the hexacoordinate metal ligand. E93 sits above the metal and solvent binding sites poised for interaction.

All structural models presented here are broadly similar to one another and to the structural model deposited to the PDB (RMSD=0.217 between *Bp*Rbr_q-apo_ and 4DI0) (SSGCID, 2012) (Table 1, SI Table 1). The existing *Bp*Rbr structure (PDB 4DI0) was solved in a different space group (P2_1_2_1_2_1_), and iron(III) is modeled into some of the di-metal binding sites. Specifically, one four-helix bundle contains two iron atoms, each modeled at 50% occupancy into both sites. The second four-helix bundle contains one iron atom modeled at 50% occupancy at site one, and no metal modeled in site two. In an effort to solve the structure of a true apo *Bp*Rbr, we treated *Bp*Rbr extensively with EDTA to chelate metals carried through the purification process. Despite these efforts, our structures of *Bp*Rbr_q-apo_ contain density for residual metal bound at the metal binding site at occupancies less than 15% (Table 2). Across the series of structures, the di-metal site consistently shows metal binding with octahedral geometry in both sites. Slight variability in the exact position of the metal ions are observed for the different structures, with the shortest distance (3.75 Å) between manganese atoms (SI Table 2). The largest variability observed is in site one coordinating residue E20 and bridging E124, which twist relative to one another and the metal ion in site one.

**Table 1.**
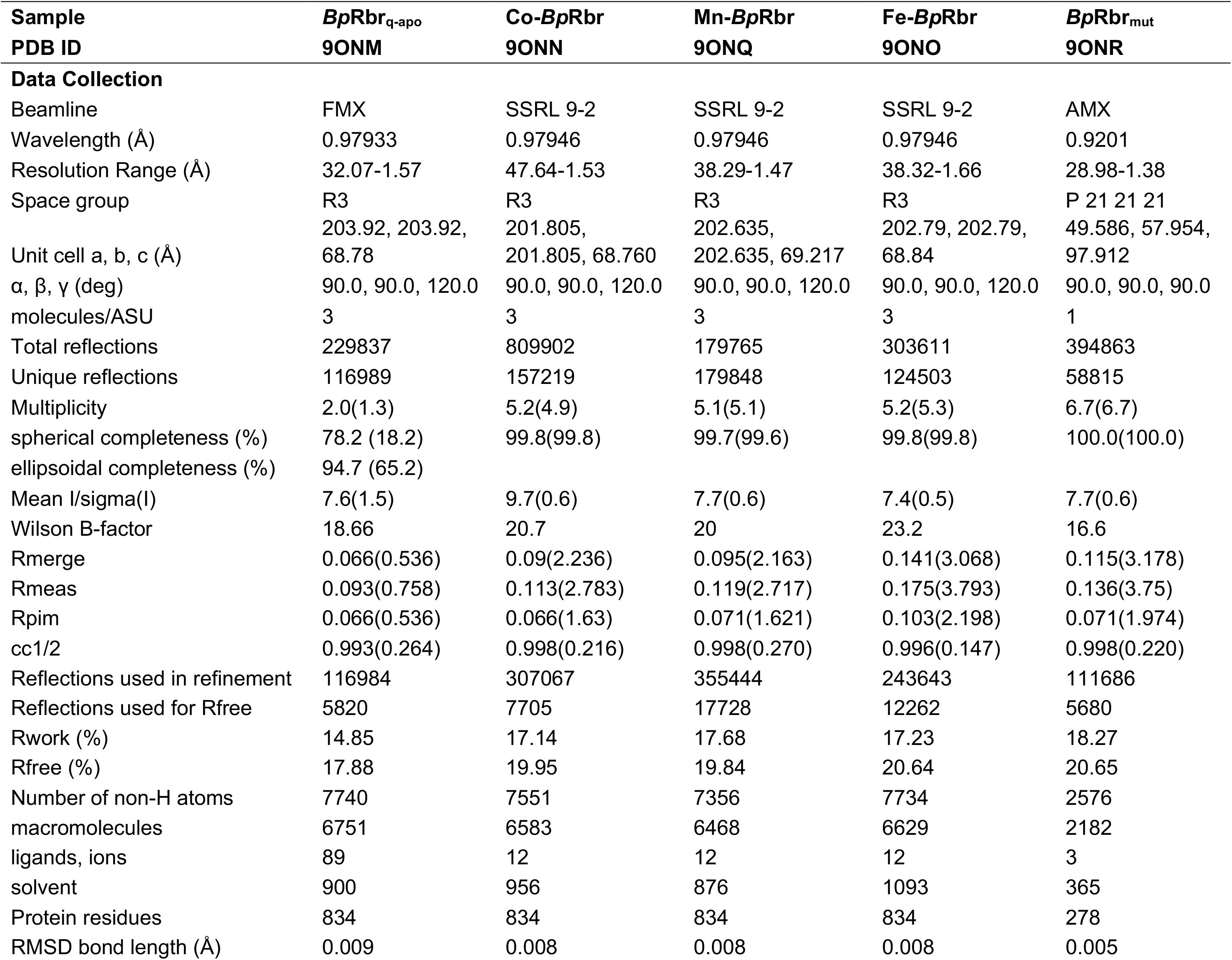
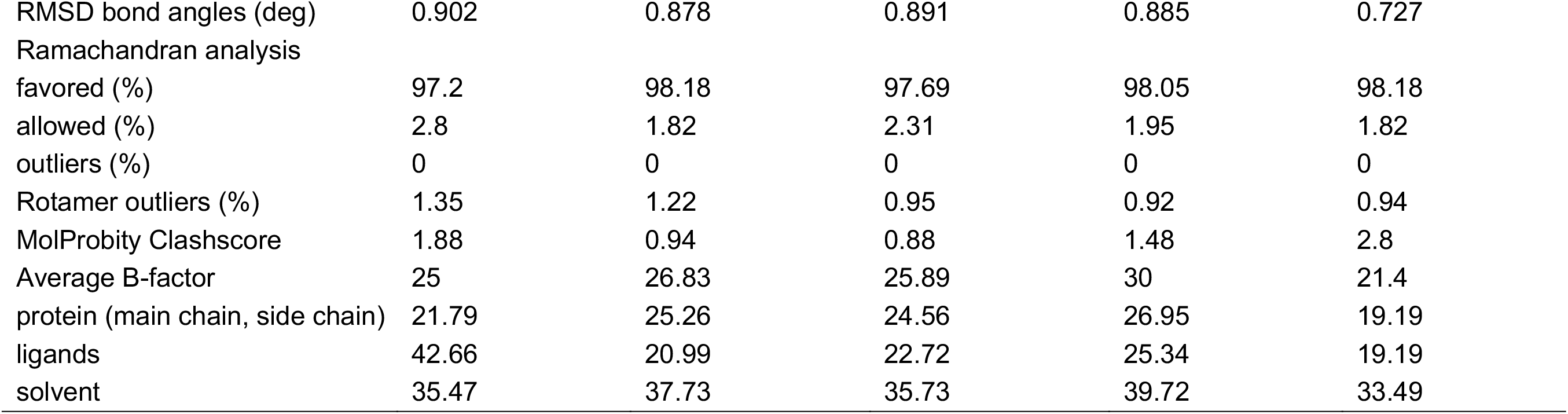
Crystallography Statistics. Values in parentheses represent inner shell values. # denotes dataset was truncated with an ellipsoidal cutoff in StarAniso (Tickle *et al*., 2018), limited reported completeness by spherical truncation methods

**Table 2.** Average refined metal occupancies for *Bp*Rbr structures.

| structure | PDB | metal modeled | average occupancy site 1 | average occupancy site 2 |
| --- | --- | --- | --- | --- |
| ***BpRbr*<sub>q-apo</sub>** | 9ONM | Fe(III) | 0.13 | 0.13 |
| **Fe-*BpRbr*** | 9ONO | Fe(III) | 0.55 | 0.41 |
| **Mn-*BpRbr*** | 9ONQ | Mn(II) | 0.22 | 0.17 |
| **Co-*BpRbr*** | 9ONN | Co(II) | 0.69 | 0.54 |

### Evidence of promiscuous metal binding in BpRbr

Although Fe(III) was modeled into the existing *Bp*Rbr structure (4DI0) (SSGCID, 2012), we sought to determine definitive structures of *Bp*Rbr bound to a series of hexacoordinate metals in order to help elucidate the physiological metal species and define metal-binding promiscuity. Di-iron in the reduced form has been identified as the active metal species for some rubrerythrins, including *Pf*Rbr and *Nv*Rbr (Weinberg *et al*., 2004, Dillard *et al*., 2011, Coulter *et al*., 1999, Barreiro *et al*., 2023). SulE is closely related to *Bp*Rbr as it exhibits the same domain-swapped conformation and lack of rubredoxin domain. In SulE, Fe(II), Mn(II), Co(II), and Ni(II) show peroxidase activity, and Mn(II), Co(II), and Ni(II) (but not iron) exhibit catalase activity (Jeoung *et al*., 2021, Lennartz *et al*., 2022). To examine metal binding in *Bp*Rbr, we solved a suite of metal-bound *Bp*Rbr structures bound to iron, manganese and cobalt (Table 1, SI Fig 1). Due to the observation that *Bp*Rbr_q-apo_ retained trace levels of metal in the crystal despite chelation treatment (SI Fig 2), we were unable to directly assign metal identities in our initial structures based on structural information alone. Furthermore, we could not say with confidence that both metal sites bound the same metal species.

To address this issue, we used EDX spectroscopy on crystals before X-ray diffraction data were collected. EDX spectroscopy provides element specific identification by measuring emission of K_α_ and K_β_ energies upon exposure to X-rays (Bowman *et al*., 2016). Surprisingly, EDX spectra on the crystals revealed that crystals contain heterogeneous combinations of metals, even when only a single metal was supplied to the protein sample at the crystallization stage, indicating unexpected heterogeneity in metal content for *Bp*Rbr crystals (Fig 3). Representative spectra are shown for two *Bp*Rbr crystals. The first crystal was grown in the presence of only manganese; despite this, iron and cobalt are observed in the sample but no manganese. The second crystal was grown in the grown in the presence of cobalt only, but both cobalt and iron are observed in the sample. We have even observed two crystals harvested from the same drop with differing amounts of iron contamination (data not shown). These results provide evidence that *Bp*Rbr is a promiscuous metal binding protein that will readily coordinate tightly to available metals as the iron was probably carried through from expression. By screening each crystal with EDX spectroscopy prior to X-ray data collection, we successfully identified crystals that contained only single, specific metals. Therefore, EDX spectra were subsequently used as a selection criterion for all X-ray data collected for structural models presented to avoid ambiguous metal assignment during structural modeling.

**Figure 3:**
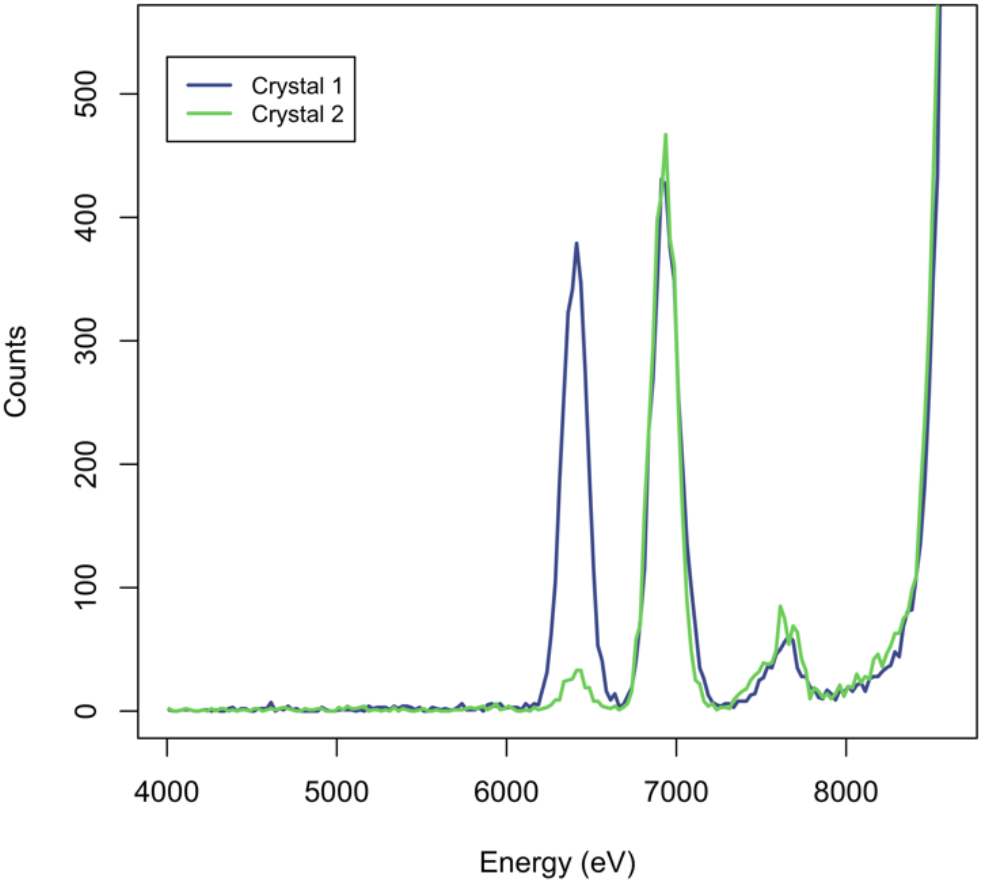
EDX spectra reveal presence of multiple metals from two different crystals. The slate blue line shows that Crystal 1 contains both iron and cobalt (iron K_a_ emission energy 6,403.84 eV and cobalt K_a_ emission energy at 6,930.32 eV). The green line shows that Crystal 2 contains primarily cobalt. EDX spectra for these samples data taken from beamline 12-2 at SSRL.

### Structures of BpRbr using coupled X-ray diffraction and EDX spectroscopy

Coupling EDX spectroscopy with X-ray crystallography enabled selection of crystals that only had one metal species in the sample, allowing for unambiguous metal assignment in the di-metal site from the resulting diffraction data. Structures were solved of *Bp*Rbr_q-apo_, Fe-, Mn-, and Co-*Bp*Rbr at 1.57, 1.66, 1.47 and 1.53 Å resolution, respectively (Table 1). In all metal-bound structural models presented here, metal identity was verified by EDX spectroscopy for the single crystal that corresponds to the diffraction data (Fig 5). Each di-metal site has one to three water molecules sitting between the di-metal site and E93 (SI Fig 3).

Metal occupancies for each metal-bound structure were refined automatically using phenix refine, as none of the metal sites were found to be reasonably modeled with full occupancies (Table 2, Fig 4). Variability is observed in metal occupancy for all structural models, and none of the metal sites are modeled with 100% metal. Based on the expected scattering factors from the metal atoms, when using the nickel k-edge excitation (8333 eV) for EDX spectroscopy, we would expect the K_α_ fluorescence intensity to follow the trend Co > Fe > Mn if the metals are present at equivalent levels. Instead, we observe the iron K_α_ as the most intense signal (intensity = 344 counts), then cobalt, then manganese (316 and 151, respectively) (Fig 4). These fluorescence intensities correlate with the modeled occupancies in the metal-bound structures (Fig 4), providing further evidence of metal-dependent binding affinity in the di-metal site. Iron and cobalt models have higher occupancy than manganese in all sites, with correspondingly smaller B-factors. For *Bp*Rbr_q-apo_ structure, the sites have 8%-20% occupancy, which we annotate as Fe(III) because of the slight presence of iron in the EDX spectrum for the crystal (SI Fig 2). Differences in water coordination at the metal binding site were also observed within different dimers of the same structures and between structures, suggestive of the ability to coordinate a substrate molecule such as peroxide (SI Fig 3).

**Figure 4:**
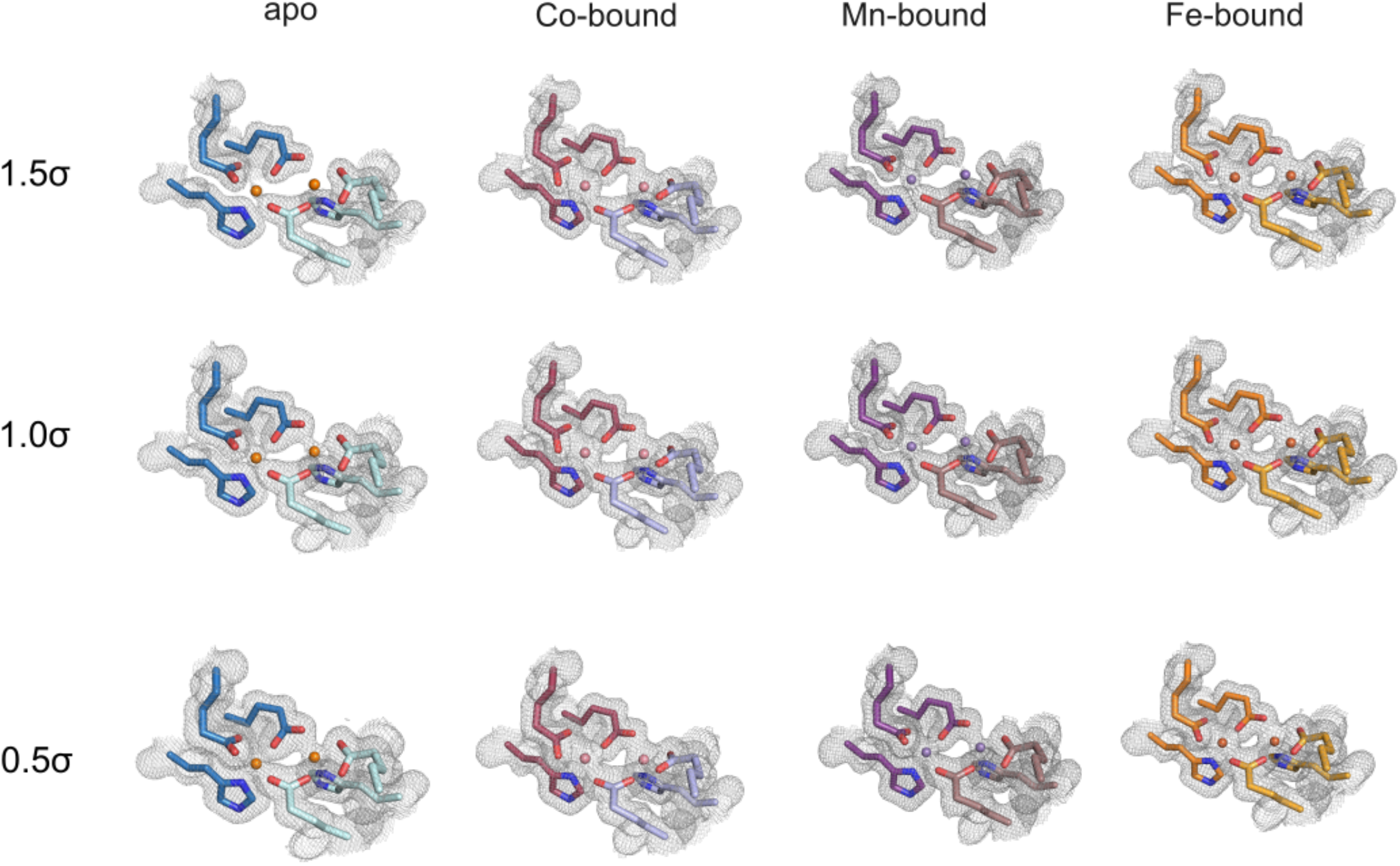
Electron density maps show differences in metal occupancy between structures. For the quasi-apo and metal-bound structures, the 2F_0_-F_c_ map is shown contoured to 1.5, 1.0 and 0.5σ. Fe(III) are shown in orange, Co(II) are shown in salmon, Mn(II) are shown in violet.

Across the series of structures, the di-metal site consistently shows metal binding with octahedral geometry in both sites (Fig 4). Slight variability in the exact position of the metal ions are observed for the different structures, with Fe- and Co-*Bp*Rbr di-metal distances closer to one another relative to the observed distances between the two metal sites in Mn-*Bp*Rbr and *Bp*Rbr_q-apo_. All metal-bound structures have very similar residue positioning in the metal-binding pocket (SI Fig 4). Asymmetry in the occupancy of site 1 and site 2 metals is observed (Fig 4, Table 2), which may be related to enzymatic mechanism. As a peroxidase, the water above the metals needs to be displaced by the peroxide substrate. For metal-soaked crystals, site 1 generally has a higher refined metal occupancy and corresponding greater signal in the 2F_o_-F_c_ map (Fig 3). This was also observed for our initial tagged structures that were grown in crystallization conditions containing metals (SI Fig 1). The asymmetry observed in metal binding in *Bp*Rbr structures is consistent with observations in SulE, in which one site has higher binding affinity relative to the second site (Jeoung *et al*., 2021, Lennartz *et al*., 2022).

### Metal binding site presented by four-helix bundle is structurally pre-formed

To determine whether metals are required for overall protein folding in *Bp*Rbr, we designed a mutant protein which substitutes the glutamate residues which bridge both metal sites (Glu53, Glu124) to alanine. In order to compare the structures of the wild-type and mutant proteins, we solved the crystallographic structure of *Bp*Rbr_mut_, which crystallized in the P2_1_2_1_2_1_ space group and diffracted to 1.38 Å with a single domain-swapped dimer in the asymmetric unit. The structure of *Bp*Rbr_mut_ aligns closely with the overall folding of the *Bp*Rbr_q-apo_ structure using Pymol alignment (RMSD = 0.550). At the active site, the metal-coordinating residues lie in relatively similar positions (RMSD = 0.301) (Fig 6A). In order to analyze electron density for metals in the metal binding pocket for *Bp*Rbr_q-apo_ and mutant structures, we generated a polder map for the iron atoms modeled into the q-apo structure. Polder maps are useful for this analysis as they are omit maps that also remove bulk solvent contributions in the vicinity of the omitted atoms. To fit metals into the mutant structure, we aligned the metal binding pocket residues for the *Bp*Rbr_q-apo_ structure and the *Bp*Rbr_mut_ structure, extracted the coordinates of the iron atoms, and incorporated the iron atoms into the mutant structure. The modeled iron atoms were then used to generate a polder map for the mutant structure in the metal binding pocket. *Bp*Rbr_mut_ exhibits no additional density within the metal-binding pocket in the polder analysis, suggesting that the E53A and E124A mutations prevent *Bp*Rbr from coordinating metal species (Fig 6B). Taken together, the relatively unchanged positions of the metal-binding pocket residues and the lack of metal density suggest that the metal-binding pocket of *Bp*Rbr is pre-formed and folds independently of coordinated metal. This conclusion is also supported by the partial occupancy of all three metal species in their respective structures.

**Figure 5:**
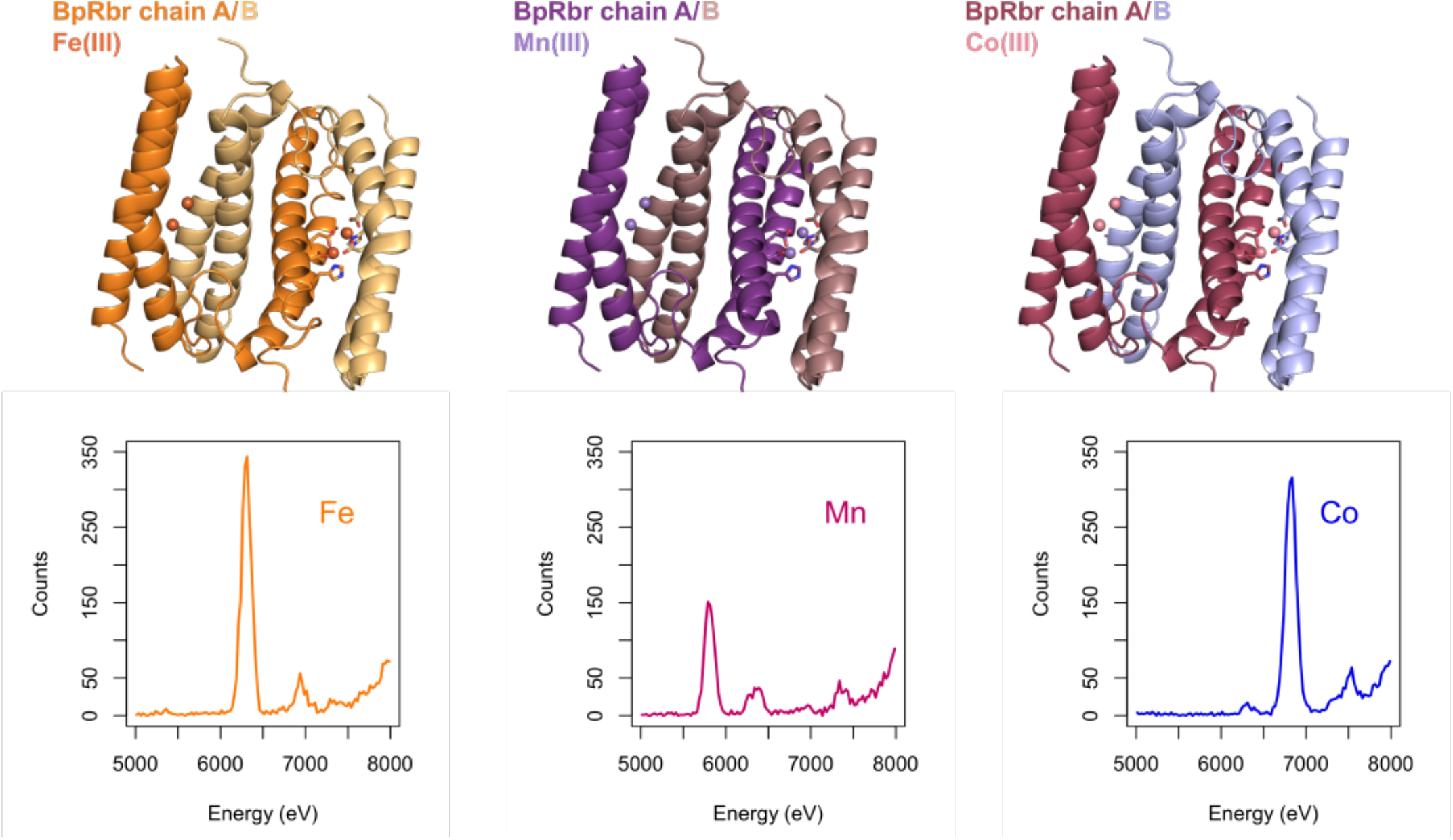
EDX spectra from single crystals corresponding to data for PDB 9ONO, 9ONQ, 9ONN. Diffraction data were collected on single crystals after first verifying presence of a single metal using EDX spectroscopy. Each EDX spectrum corresponds to the same sample on which diffraction data were collected and structures solved. EDX spectra and diffraction data for these samples collected on SSRL beamline 9-2.

**Figure 6:**
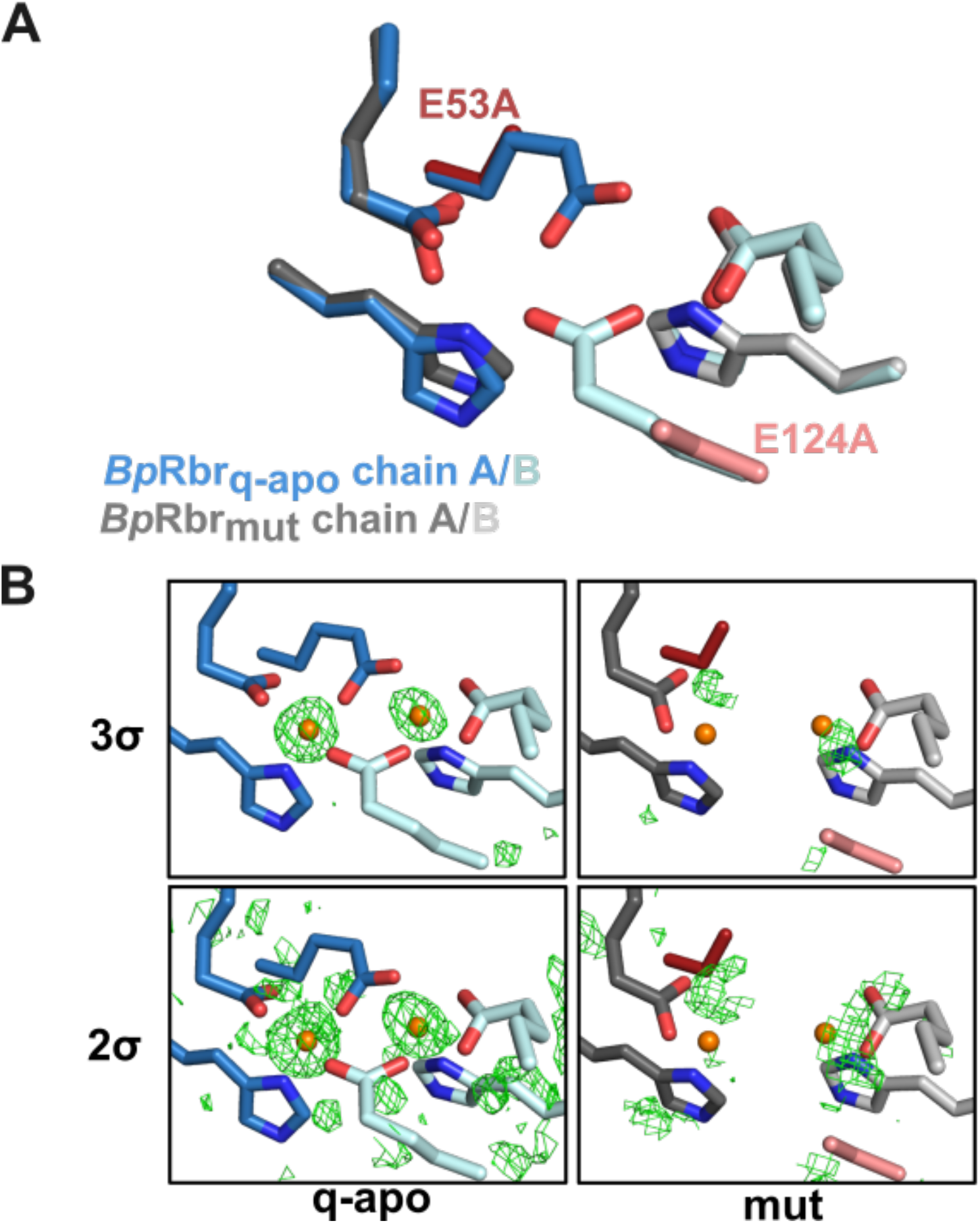
Metal binding site is pre-formed in *Bp*Rbr four helix bundles. A) Overlay of *Bp*Rbr_q-apo_ metal binding pocket with *Bp*Rbr_mut_ binding pocket show a high degree of similarity in residue position. (RMSD = 0.550). *Bp*Rbr_q-apo_ chain A is represented in dark blue and chain B is represented in teal. *Bp*Rbr_mut_ chain A is represented in dark grey and chain B is represented in light grey. Mutations E53A and E124A are rendered in firebrick red and salmon, respectively. B) Polder maps of binding pocket metals in *Bp*Rbr_q-apo_ and *Bp*Rbr_mut_ contoured to 3σ and 2σ. Modeled Fe(III) atoms are colored in orange and Polder difference maps are represented in green mesh.

## Conclusions

Rubrerythrin is an enzyme involved in oxidative stress response in bacteria. *Bp*Rbr is speculated to be a reductase with peroxide as its native substrate. It shows atypical structural features compared to Rbr from other organisms. We observe that the structure is composed of a domain swapped dimer and is missing the canonical rubredoxin domain found in Rbr structural models from other organisms. These structural features speak to the potential role of oxygen exposure in the evolution of multiple Rbr forms from different organisms. Given the key role played by rubrerythrin proteins in OSR in a wide variety of microorganisms, this important class of enzymes has already been probed from an evolutionary perspective (Cardenas *et al*., 2016). Cardenas and coworkers used protein sequence-based phylogenomics to identify multiple subgroups within the Rbr family. The largest subgroup is the ‘classical’ form, of which ∼99% of the sequences contain the rubredoxin domain. *Bp*Rbr is a member of a smaller subgroup, the ‘aerobic-type’, which exclusively contains Rbrs lacking the rubredoxin domain. Most of the organisms in this group are facultative aerobes from a diverse array of bacterial and archaeal taxa. Further, many of the other ‘aerobic-type’ species found to contain an Rbr with no rubredoxin domain are extremophiles (Sato *et al*., 2012, Jeoung *et al*., 2021). The rubredoxin domain is mechanistically required in the ‘classical’ Rbr peroxidase activity, as an electron transfer partner important in the reduction of peroxide. Further work is required to determine whether reduced metals will alter the *Bp*Rbr metal binding site. In other metalated Rbr enzymes, active metals for catalysis are in the reduced oxidation state, while we have modeled iron in the oxidized state. We chose to do so because crystallization sample was prepared under aerobic conditions, and it is therefore likely that the iron is oxidized. It is also likely that at least in the case of the iron-bound species, an anaerobic chamber will be required to generate crystals bound to ferrous iron. EDX spectroscopy cannot distinguish between metal oxidation states, but does provide information about what metal is present in the sample. X-ray Absorption Near Edge Structure (XANES) or X-ray emission spectroscopy (XES) are methods that can be used to validate metal oxidation states. However, in Rbr these approaches are complicated by the presence of two asymmetric metals per binding site, which may benefit from more stringent approaches to local ionization state determination like spatially resolved anomalous dispersion (SpReAD) analysis (Einsle *et al*., 2007, Lennartz *et al*., 2022).

Our work uncovering metal promiscuity and the pre-formed metal-binding site in *Bp*Rbr speaks to the potential role of adaptation to an oxygen containing environment as crucial for the structural evolution of *Bp*Rbr. We hypothesize that the other members of the rubrerythrin operon in *Bp* may be the electron transfer partner that replaces the rubredoxin domain found in the ‘classical’ Rbrs. The structural work raises new questions about metabolic flexibility in Rbrs from aerobic and extremophilic organisms; structures with domain swapped dimers forming the metal-binding four-helix bundle have thus far been found only in aerobic and extremophile environments. These results highlight the need for further investigation into these non-classical systems. In addition, our work has demonstrated the importance of rigorously assessing metal identity in samples, given the promiscuity that we observe in metal-binding in *Bp*Rbr. We have used *in crystallo* EDX spectroscopy as a selection criteria for structural studies. The ideal solution to examine metal speciation in metalloproteins would be to collect diffraction data both above and below the anomalous absorption edge (Handing *et al*., 2018). However, in samples that have heterogeneity in metal speciation, as we have observed for *Bp*Rbr, that is challenging. The challenge arises first because of the sheer number of datasets that would need to be collected (above and below each metal absorption edge). Second, the *Bp*Rbr crystals are sensitive to radiation damage, and so the X-ray exposure dose must be carefully monitored. In samples sensitive to radiation damage, which includes many metalloproteins, it can be difficult to collect sufficient data on individual crystals to obtain the anomalous data required. Here we present a way to use the correlation between the EDX intensity and the modeled metal occupancy to provide an additional, lower radiation dose tool for probing aspects of metal-binding in these systems. We believe that the use of EDX spectroscopy expands our ability to measure metal speciation in metalloprotein systems.

## Abbreviations

(*Bp*): *Burkholderia pseudomallei*
(*Pf*): *Pyroccocus furiosus*
(*Nv*): *Nitratidesulfovibrio vulgaris*
(Rbr): Rubrerythrin
(SulE): sulerythrin
(*Bp*Rbr_q-apo_): quasi-apo
(*Bp*Rbr_mut_): *Bp*Rbr-E53A/E124A mutant
(EDX): Energy dispersive X-ray spectroscopy
(OSR): oxidative stress response
(DUF): domain of unknown function

## Acknowledgements and Funding

This research was supported in part by the Intramural Research Program of the NIH (DCFM) and in part by the Dr. Louis Skarlow Memorial Fund (SEJB).

Initial crystallization conditions for rubrerythrin crystals were determined with high-throughput screening at the National Crystallization Center at UB HWI supported by NIH NIGMS R24 National Resource Grant (R24GM141256). The Bowman Lab is a Member of the SBGrid Consortium, and structural biology applications used in this project were compiled and configured by SBGrid (Morin *et al*., 2013). Phenix software use is supported by NIH NIGMS R24 National Resource Grant (R24GM141254).

This research used beamlines 9-2 and 12-2 at Stanford Synchrotron Radiation Lightsource. Use of the Stanford Synchrotron Radiation Lightsource, SLAC National Accelerator Laboratory, is supported by the U.S. Department of Energy, Office of Science, Office of Basic Energy Sciences under Contract No. DE-AC02-76SF00515. The SSRL Structural Molecular Biology Program is supported by the DOE Office of Biological and Environmental Research, and by the National Institutes of Health, National Institute of General Medical Sciences (P30GM133894). The contents of this publication are solely the responsibility of the authors and do not necessarily represent the official views of NIGMS or NIH.

This research used beamlines 17-ID-1 and 17-ID-2 of the National Synchrotron Light Source II, a U.S. Department of Energy (DOE) Office of Science User Facility operated for the DOE Office of Science by Brookhaven National Laboratory under Contract No. DE-SC0012704. The Center for BioMolecular Structure (CBMS) is primarily supported by the National Institutes of Health, National Institute of General Medical Sciences (NIGMS) through a Center Core P30 Grant (P30GM133893), and by the DOE Office of Biological and Environmental Research (KP1607011).

The contents of this publication are solely the responsibility of the authors and do not necessarily represent the official views of NIGMS or NIH.

## CrediT author statement

**GR Budziszewski**: Investigation, Data Curation, Formal Analysis, Writing – Original Draft, Writing – Review & Editing, Visualization **ML Lynch** Formal Analysis, Software, Writing – Original Draft, Writing – Review & Editing, Visualization **ME Snell** Investigation, Data Curation, Writing – Review & Editing, **DCF Monteiro** Conceptualization, Investigation, Data Curation, Formal Analysis, Writing – Original Draft, Writing – Review & Editing, Visualization, Funding Acquisition **SEJ Bowman** Conceptualization, Investigation, Data Curation, Formal Analysis, Writing – Original Draft, Writing – Review & Editing, Visualization, Funding Acquisition

## Notes

### Competing Interest Statement

The authors have declared no competing interest.

## References

Barreiro, D. S., Oliveira, R. N. S. & Pauleta, S. R. (2023). Bacterial Peroxidases: Multivalent Enzymes that Enable the Use of Hydrogen Peroxide for Microaerobic and Anaerobic Proliferation, Coordination Chemistry Reviews 485, 215114.

Bowman, S. E. J., Bridwell-Rabb, J. & Drennan, C. L. (2016). Metalloprotein Crystallography: More than a Structure, Accounts of Chemical Research 49, 695–702.

Budziszewski, G. R., Snell, M. E., Wright, T. R., Lynch, M. L. & Bowman, S. E. J. (2023). High-Throughput Screening to Obtain Crystal Hits for Protein Crystallography, JoVE, e65211.

Burkholderia Genome DB (2024). https://www.burkholderia.com/.

Butt, A. T. & Thomas, M. S. (2017). Iron Acquisition Mechanisms and Their Role in the Virulence of Burkholderia Species, Frontiers in Cellular and Infection Microbiology Volume 7 - 2017.

Cardenas, J. P., Quatrini, R. & Holmes, D. S. (2016). Aerobic Lineage of the Oxidative Stress Response Protein Rubrerythrin Emerged in an Ancient Microaerobic, (Hyper)Thermophilic Environment, Frontiers in Microbiology Volume 7 - 2016.

Casadevall, A. (2020). Climate Change Brings the Specter of New Infectious Diseases, The Journal of Clinical Investigation 130, 553–555.

Coulter, E. D., Shenvi, N. V. & Kurtz, D. M. (1999). NADH Peroxidase Activity of Rubrerythrin, Biochemical and Biophysical Research Communications 255, 317–323.

Dawan, J. & Ahn, J. (2022). Bacterial Stress Responses as Potential Targets in Overcoming Antibiotic Resistance, Microorganisms 10, 1385.

Dillard, B. D., Demick, J. M., Adams, M. W. W. & Lanzilotta, W. N. (2011). A Cryo-crystallographic Time Course for Peroxide Reduction by Rubrerythrin from Pyrococcus furiosus, JBIC Journal of Biological Inorganic Chemistry 16, 949–959.

Dolecek, C., Shakoor, S., Basnyat, B., Okwor, T. & Sartorius, B. (2022). Drug-resistant Bacterial Infections: We Need Urgent Action and Investment That Focus on the Weakest Link, PLOS Biology 20, e3001903.

Einsle, O., Andrade, S. L. A., Dobbek, H., Meyer, J. & Rees, D. C. (2007). Assignment of Individual Metal Redox States in a Metalloprotein by Crystallographic Refinement at Multiple X-ray Wavelengths, Journal of the American Chemical Society 129, 2210–2211.

Evans, P. (2006). Scaling and Assessment of Data Quality, Acta Crystallographica Section D 62, 72–82.

Evans, P. R. & Murshudov, G. N. (2013). How Good are My Data and What is the Resolution?, Acta Crystallographica Section D 69, 1204–1214.

Garnier, S., Ross, N., Rudis, R., Camargo, P. A., Sciaini, M. & Scherer, C. (2024). viridis(Lite) - Colorblind-Friendly Color Maps for R., doi:10.5281/zenodo.4679423, viridis package version 0.6.5, https://sjmgarnier.github.io/viridis/.

Handing, K. B., Niedzialkowska, E., Shabalin, I. G., Kuhn, M. L., Zheng, H. & Minor, W. (2018). Characterizing metal-binding sites in proteins with X-ray crystallography, Nature Protocols 13, 1062–1090.

Holden, M. T. G., Titball, R. W., Peacock, S. J., Cerdeño-Tárraga, A. M., Atkins, T., Crossman, L. C., Pitt, T., Churcher, C., Mungall, K., Bentley, S. D., Sebaihia, M., Thomson, N. R., Bason, N., Beacham, I. R., Brooks, K., Brown, K. A., Brown, N. F., Challis, G. L., Cherevach, I., Chillingworth, T., Cronin, A., Crossett, B., Davis, P., DeShazer, D., Feltwell, T., Fraser, A., Hance, Z., Hauser, H., Holroyd, S., Jagels, K., Keith, K. E., Maddison, M., Moule, S., Price, C., Quail, M. A., Rabbinowitsch, E., Rutherford, K., Sanders, M., Simmonds, M., Songsivilai, S., Stevens, K., Tumapa, S., Vesaratchavest, M., Whitehead, S., Yeats, C., Barrell, B. G., Oyston, P. C. F. & Parkhill, J. (2004). Genomic Plasticity of the Causative Agent of Melioidosis, Burkholderia pseudomallei, Proceedings of the National Academy of Sciences 101, 14240–14245.

Jeoung, J.-H., Rünger, S., Haumann, M., Neumann, B., Klemke, F., Davis, V., Fischer, A., Dau, H., Wollenberger, U. & Dobbek, H. (2021). Bimetallic Mn, Fe, Co, and Ni Sites in a Four-Helix Bundle Protein: Metal Binding, Structure, and Peroxide Activation, Inorganic Chemistry 60, 17498–17508.

Jitprasutwit, S., Ong, C., Juntawieng, N., Ooi, W. F., Hemsley, C. M., Vattanaviboon, P., Titball, R. W., Tan, P. & Korbsrisate, S. (2014). Transcriptional Profiles of Burkholderia pseudomallei Reveal the Direct and Indirect Roles of Sigma E under Oxidative Stress Conditions, BMC Genomics 15, 787.

Kabsch, W. (2010). XDS, Acta Crystallographica Section D 66, 125–132.

Lennartz, F., Jeoung, J.-H., Ruenger, S., Dobbek, H. & Weiss, M. S. (2022). Determining the Oxidation State of Elements by X-ray Crystallography, Acta Crystallographica Section D 78, 238–247.

Liebschner, D., Afonine, P. V., Baker, M. L., Bunkoczi, G., Chen, V. B., Croll, T. I., Hintze, B., Hung, L.-W., Jain, S., McCoy, A. J., Moriarty, N. W., Oeffner, R. D., Poon, B. K., Prisant, M. G., Read, R. J., Richardson, J. S., Richardson, D. C., Sammito, M. D., Sobolev, O. V., Stockwell, D. H., Terwilliger, T. C., Urzhumtsev, A. G., Videau, L. L., Williams, C. J. & Adams, P. D. (2019). Macromolecular Structure Determination Using X-rays, Neutrons and Electrons: Recent Developments in Phenix, Acta Crystallographica Section D 75, 861–877.

Liebschner, D., Afonine, P. V., Moriarty, N. W., Poon, B. K., Sobolev, O. V., Terwilliger, T. C. & Adams, P. D. (2017). Polder Maps: Improving OMIT Maps by Excluding Bulk Solvent, Acta Crystallographica Section D 73, 148–157.

Lynch, M. L., Snell, M. E., Potter, S. A., Snell, E. H. & Bowman, S. E. J. (2023). 20 Years of Crystal Hits: Progress and Promise in Ultrahigh-throughput Crystallization Screening, Acta Crystallographica Section D 79, 198–205.

Mora, C., McKenzie, T., Gaw, I. M., Dean, J. M., von Hammerstein, H., Knudson, T. A., Setter, R. O., Smith, C. Z., Webster, K. M. & Patz, J. A. (2022). Over Half of Known Human Pathogenic Diseases Can Be Aggravated by Climate Change, Nature climate change 12, 869–875.

Morin, A., Eisenbraun, B., Key, J., Sanschagrin, P. C., Timony, M. A., Ottaviano, M. & Sliz, P. (2013). Collaboration Gets the Most out of Software, eLife 2, e01456.

Murshudov, G. N., Skubak, P., Lebedev, A. A., Pannu, N. S., Steiner, R. A., Nicholls, R. A., Winn, M. D., Long, F. & Vagin, A. A. (2011). REFMAC5 for the Refinement of Macromolecular Crystal Structures, Acta Crystallographica Section D 67, 355–367.

Petras, J. K., Elrod, M. G., Ty, M. C., Dawson, P., O’Laughlin, K., Gee, J. E., Hanson, J., Boutwell, C., Ainsworth, G., Beesley, C. A., Saile, E., Tiller, R., Gulvik, C. A., Ware, D., Sokol, T., Balsamo, G., Taylor, K., Salzer, J. S., Bower, W. A., Weiner, Z. P., Negrón, M. E., Hoffmaster, A. R. & Byers, P. (2023). Locally Acquired Melioidosis Linked to Environment — Mississippi, 2020–2023, New England Journal of Medicine 389, 2355–2362.

Phillips, E. D. & Garcia, E. C. (2024). Burkholderia pseudomallei, Trends in Microbiology 32, 105–106.

R Core Team (2024). R: A Language and Environment for Statistical Computing, https://www.R-project.org/.

Sato, Y., Kameya, M., Fushinobu, S., Wakagi, T., Arai, H., Ishii, M. & Igarashi, Y. (2012). A Novel Enzymatic System against Oxidative Stress in the Thermophilic Hydrogen-Oxidizing Bacterium Hydrogenobacter thermophilus, PLOS ONE 7, e34825.

Schweizer, H. P. (2012). Mechanisms of Antibiotic Resistance in Burkholderia pseudomallei: Implications for Treatment of Melioidosis, Future Microbiology 7, 1389–1399.

Soltis, S. M., Cohen, A. E., Deacon, A., Eriksson, T., Gonzalez, A., McPhillips, S., Chui, H., Dunten, P., Hollenbeck, M., Mathews, I., Miller, M., Moorhead, P., Phizackerley, R. P., Smith, C., Song, J., van dem Bedem, H., Ellis, P., Kuhn, P., McPhillips, T., Sauter, N., Sharp, K., Tsyba, I. & Wolf, G. (2008). New Paradigm for Macromolecular Crystallography Experiments at SSRL: Automated Crystal Screening and Remote Data Collection, Acta Crystallographica Section D 64, 1210–1221.

SSGCID (2012). The structure of Rubrerythrin from Burkholderia pseudomallei., 10.2210/pdb4di0/pdb.

Sztukowska, M., Bugno, M., Potempa, J., Travis, J. & Kurtz Jr, D. M. (2002). Role of Rubrerythrin in the Oxidative Stress rRsponse of Porphyromonas gingivalis, Molecular microbiology 44, 479–488.

Taboada, B., Estrada, K., Ciria, R. & Merino, E. (2018). Operon-mapper: A Web Server for Precise Operon Identification in Bacterial and Archaeal Genomes, Bioinformatics 34, 4118–4120.

Thompson, A., Attwood, D., Gullickson, E., Howells, M., Kim, K.-J., Kirz, J., Kortright, J., Lindau, I., Liu, Y., Pianetta, P., Robinson, A., Scofield, J., Underwood, J., Williams, G. & Winick, H. (2009). X-ray Data Booklet, https://xdb.lbl.gov/Section1/Table_1-2.pdf.

Tickle, I., Flensburg, C., Keller, P., Paciorek, W., Sharff, A., Vonrhein, C. & Bricogne, G. (2018). STARANISO.

Torres, A. G. (2023). The Public Health Significance of Finding Autochthonous Melioidosis Cases in the Continental United States, PLOS Neglected Tropical Diseases 17, e0011550.

Vagin, A. & Teplyakov, A. (1997). MOLREP: an Automated Program for Molecular Replacement, Journal of Applied Crystallography 30, 1022–1025.

Vatansever, F., de Melo, W. C. M. A., Avci, P., Vecchio, D., Sadasivam, M., Gupta, A., Chandran, R., Karimi, M., Parizotto, N. A., Yin, R., Tegos, G. P. & Hamblin, M. R. (2013). Antimicrobial Strategies Centered Around Reactive Oxygen Species – Bactericidal Antibiotics, Photodynamic Therapy, and Beyond, FEMS Microbiology Reviews 37, 955–989.

Vonrhein, C., Flensburg, C., Keller, P., Sharff, A., Smart, O., Paciorek, W., Womack, T. & Bricogne, G. (2011). Data Processing and Analysis with the autoPROC Toolbox, Acta Crystallographica Section D 67, 293–302.

Weinberg, M. V., Jenney, F. E., Cui, X. & Adams, M. W. W. (2004). Rubrerythrin from the Hyperthermophilic Archaeon Pyrococcus furiosus Is a Rubredoxin-Dependent, Iron-Containing Peroxidase, Journal of Bacteriology 186, 7888–7895.

Winn, M. D., Ballard, C. C., Cowtan, K. D., Dodson, E. J., Emsley, P., Evans, P. R., Keegan, R. M., Krissinel, E. B., Leslie, A. G. W., McCoy, A., McNicholas, S. J., Murshudov, G. N., Pannu, N. S., Potterton, E. A., Powell, H. R., Read, R. J., Vagin, A. & Wilson, K. S. (2011). Overview of the CCP4 Suite and Current Developments, Acta Crystallographica Section D 67, 235–242.

Woods, D. & Nierman, W. (2005). Burkholderia pseudomallei 1710b chromosome II, complete sequence, https://www.ncbi.nlm.nih.gov/nuccore/CP000125.

