## SupplementalInformation for "*Burkholderia pseudomallei* rubrerythrin promiscuously binds metals in a structurally pre-formed bimetallic binding site"

<u>Article title</u>

### Supplemental Information

Three structures for *Bp*Rbr were initially determined, using the construct that contained the His-tag. His-tagged 'apo'-*Bp*Rbr (PDB 8FUH), 'Mn'-*Bp*Rbr (PDB 8FXD), and 'Fe'-*Bp*Rbr (PDB 8FVV); we use the 'apostrophe' notation because we cannot be certain of the metal identity for these structures, as the crystal samples exhibited significant metal heterogeneity. We include these structures in the Supplemental so that experimental details are available.

<u>Data collection and structure solution for His-tagged *Bp*Rbr</u>

Diffraction data were collected at NSLS-II at the AMX beamline 17-ID-1 (4.3 x $10^{12}$ ph/s, 0.920100 Å, 100 K) using an Eiger 9M detector for the His-tagged 'apo'-*Bp*Rbr crystals and at the FMX beamline 17-ID-2 (4 x $10^{12}$ ph/s, 0.97933 Å, 100K) using a Dectris Pilatus 6M detector for the His-tagged 'Mn'- and 'Fe'-*Bp*Rbr crystals.

Initial data reduction was performed using the automated pipeline available at NSLS-II, using autoproc (Vonrhein *et al.*, 2011). For the apo dataset, the output from the integration step from autoproc was scaled and merged using Aimless (Evans & Murshudov, 2013), phased using molecular replacement using PDB 4DI0 (SSGCID, 2012). The model was refined using Refmac5 and re-built manually using Coot (Murshudov *et al.*, 2011). The tagged 'apo' *Bp*Rbr model (PDB 8FUH) was then used as a starting model for the molecular replacement of the tagged 'Mn'-*Bp*Rbr and 'Fe'-*Bp*Rbr structures (PDB 8FXD and 8FVV, respectively). Tagged 'Mn'-*Bp*Rbr and 'Fe'-*Bp*Rbr data were integrated with the autoproc pipeline with no resolution cut-off, the unscaled, unmerged file used to determine the space group using Pointless (CCP4I2, part of Aimless) and the output scaled and merged and resolution cutoff determined anisotropically using the StarAniso server (Tickle *et al.*, 2018). The merged .mtz files from StarAniso were then used for structure solution by direct refinement against the tagged 'apo' *Bp*Rbr structure and manually rebuilt using Coot. The metal and metal-coordinating water occupancies were refined using phenix refine (Liebschner *et al.*, 2019).

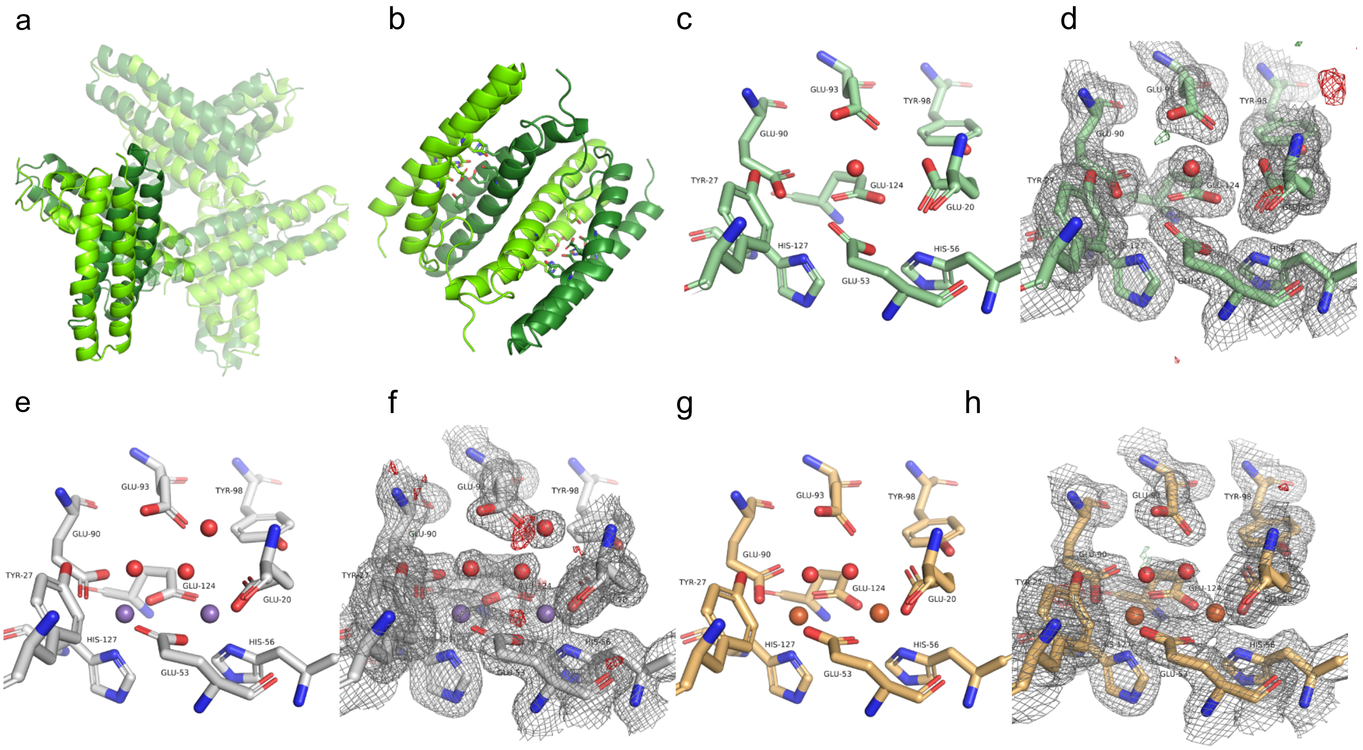


**SI Figure 1: Structures of His-tagged Rbr samples.** a) Asymmetric unit cell for His-tagged ‘apo’ *Bp*Rbr (PDB 8FUH) in green and dark green. b) Dimer is domain swapped. c-d) Metal binding site for His-tagged Apo *Bp*Rbr. e-f) Metal binding site for His-tagged ‘Mn’-*Bp*Rbr (PDB 8FXD). g-h) Metal binding site for His-tagged ‘Fe’-*Bp*Rbr (PDB 8FVV). While metal ions are modeled into these structures, these data were collected prior to the realization that there was heterogenous and promiscuous metal binding, so we cannot be *certain* that the metal identity is correct.

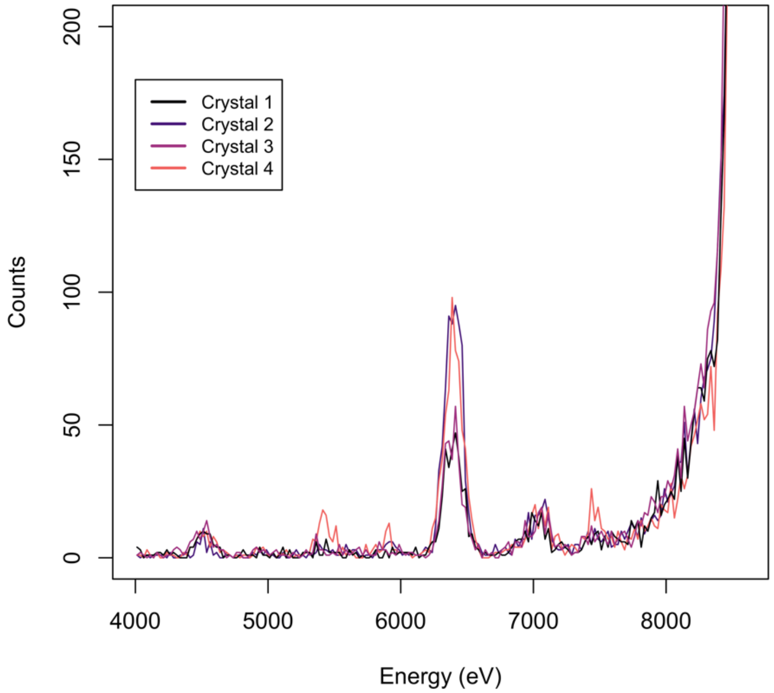


**SI Figure 2:** EDX spectra from four different Apo *Bp*Rbr crystals reveal trace amount of iron remain in the samples even after chelation. The amount of iron is lower than that observed in the metal-bound *Bp*Rbr structures, as evidenced by the lower count value for the EDX signal at the iron $K_{\alpha}$ emission energy (6,403.84 eV) (Fig 5). EDX spectra for these samples data taken from beamline 12-2 at SSRL.

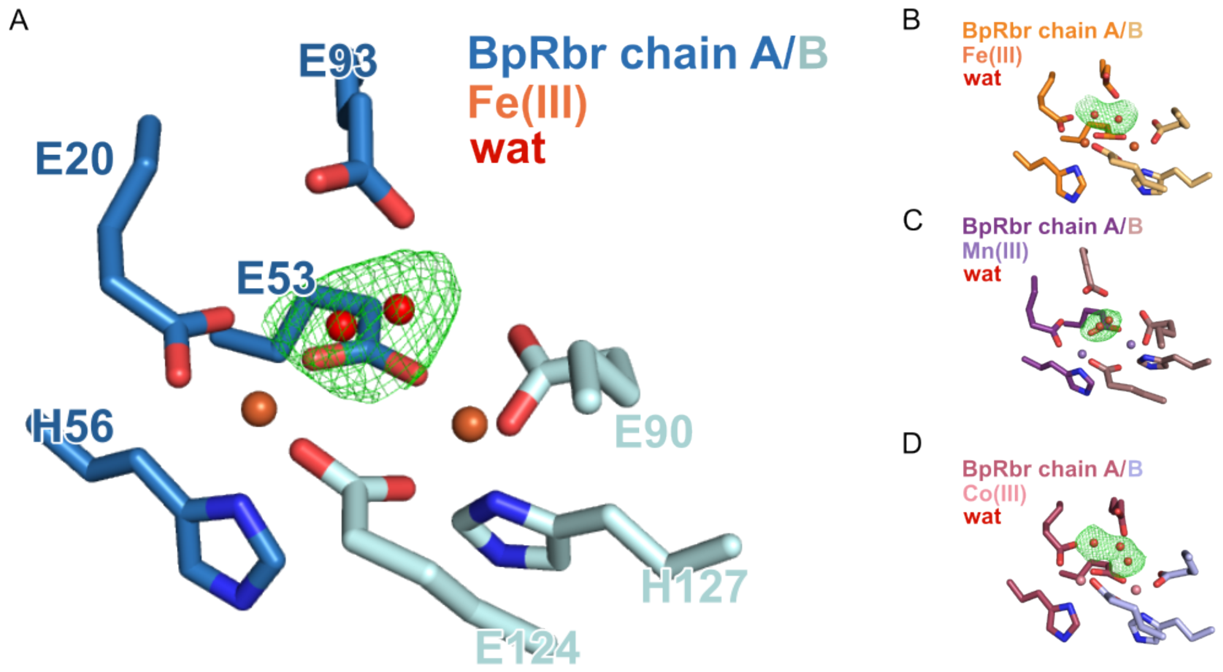


**SI Figure 3**. Polder analysis indicates the presence of mobile solvent molecules above the metal binding site consistent with the position of a peroxide substrate. Polder map is shown in green mesh and contoured to 3σ. A) Representative *Bp*Rbr$_{q\text{-}apo}$ metal binding site with a single solvent water molecule modeled in split positions with 50% occupancy each. B) Representative *Bp*Rbr-Fe metal binding site with a single solvent water molecule modeled in split positions with 50% occupancy each. C) Representative *Bp*Rbr-Mn metal binding site with a single solvent water molecule modeled in split positions with 50% occupancy each. D) Representative *Bp*Rbr-Co metal binding site with two solvent water molecules, one modeled in split positions with 50% occupancy each.

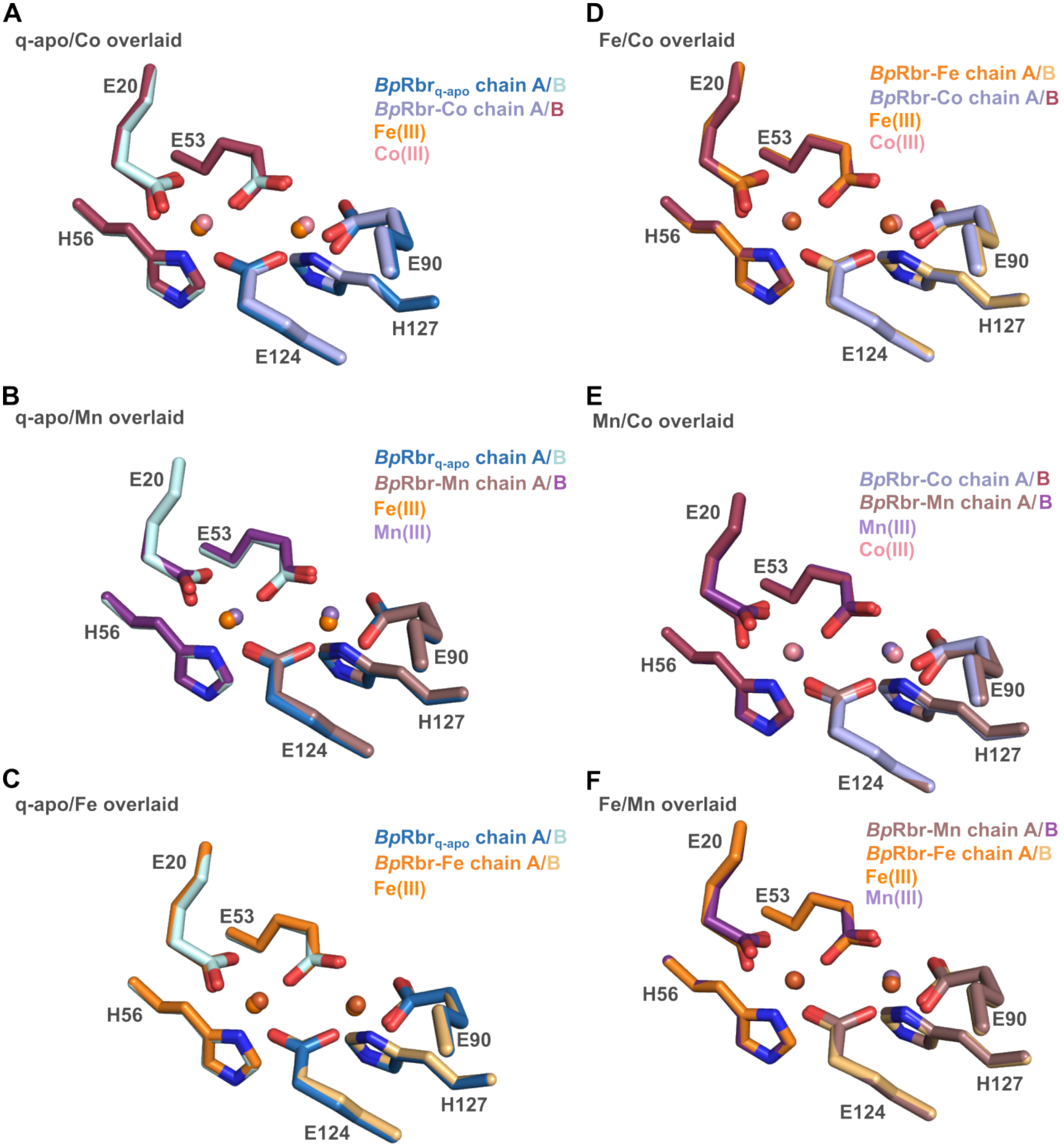


**SI Figure 4: Overlay of structures show that all metal-bound structures have very similar residue positioning in the metal-binding pocket**.

| PDB ID | His tagged 'apo'-*Bp*Rbr **8FUH** | His tagged 'Mn'-*Bp*Rbr **8FXD** | His tagged 'Fe'-*Bp*Rbr **8FVV** |
|---|---|---|---|
| Space group | R3 | R3 | R3 |
| a, b, c | 202.4, 202.4, 70.3 | 201.9, 201.9, 69.0 | 202.2, 202.2, 69.8 |
| a, b, g | 90, 90, 120 | 90, 90, 120 | 90, 90, 120 |
| Resolution range | 29.21-1.85 (1.88-1.85) | 29.14-1.58 (1.69-1.58) | 29.90-1.93 (2.07-1.93) |
| Total reflections | 323422 (14095) | 561571 (17006) | 661896 (38503) |
| unique | 90867 (4220) | 108953 (17006) | 61079 (3595) |
| completeness | 99.4 (93.4) | 93.7 (59.3) | 92.4 (61.4) |
| redundancy | 3.6 (3.3) | 5.2 (3.1) | 10.8 (10.7) |
| I/sigI (merged | 6.5 (1.4) | 9.5 (1.4) | 8.9 (1.5) |
| CC1/2 | 0.993 (0.608) | 0.998 (0.457) | 0.995 (0.597) |
| Rpim | 0.093 (0.643) | 0.050 (0.597) | 0.073 (0.607) |
| Rmeas | 0.138 (0.934) | 0.115 (1.073) | 0.241 (1.993) |
| B-factor | 14.37 | 19.83 | 25.69 |
| REFINEMENT | | | |
| Working set | 86407 | 103483 | 58081 |
| Test set | 4443 | 5467 | 2998 |
| R | 15.0 | 14.8 | 16.1 |
| Rfree | 18.8 | 17.5 | 20.9 |
| Total non-H atoms | 7498 | 7475 | 7279 |
| Ions | 0 | 12 | 12 |
| ligands | 14 | 42 | 0 |
| waters | 959 | 898 | 803 |
| Bond RMS | 0.0081 | 0.0103 | 0.0087 |
| Angle RMS | 1.42 | 1.54 | 1.45 |
| Bfactor - total | 31.87 | 22.23 | 28.10 |
| ions | 0 | 12.82 | 41.61 |
| ligands | 65.25 | 47.56 | 0 |
| waters | 35.77 | 32.24 | 35.92 |
| Favored (%) | 97.4 | 97.7 | 97.3 |
| Outliers (%) | 0.62 | 0.73 | 0.61 |

**SI Table 1**: Crystallography Statistics. Values in parentheses represent inner shell values.

| metal | distance (Å) |
|---|---|
| Fe-Fe | 3.83 |
| Co-Co | 3.88 |
| Mn-Mn | 3.75 |

**SI Table 2**: Average distance between metals in three metal-soaked *Bp*Rbr structures (PDB 9ONO, 9ONN, and 9ONQ)

**Supplemental References**


Evans, P. R. & Murshudov, G. N. (2013). How Good are My Data and What is the Resolution? ,*Acta Crystallographica Section D* **69**, 1204-1214.

Liebschner, D., Afonine, P. V., Baker, M. L., Bunkoczi, G., Chen, V. B., Croll, T. I., Hintze, B., Hung, L.-W., Jain, S., McCoy, A. J., Moriarty, N. W., Oeffner, R. D., Poon, B. K., Prisant, M. G., Read, R. J., Richardson, J. S., Richardson, D. C., Sammito, M. D., Sobolev, O. V., Stockwell, D. H., Terwilliger, T. C., Urzhumtsev, A. G., Videau, L. L., Williams, C. J. & Adams, P. D. (2019). Macromolecular Structure Determination Using X-rays, Neutrons and Electrons: Recent Developments in Phenix,*Acta Crystallographica Section D* **75**, 861-877.

Murshudov, G. N., Skubak, P., Lebedev, A. A., Pannu, N. S., Steiner, R. A., Nicholls, R. A., Winn, M. D., Long, F. & Vagin, A. A. (2011). REFMAC5 for the Refinement of Macromolecular Crystal Structures,*Acta Crystallographica Section D* **67**, 355-367.

SSGCID (2012). *The structure of Rubrerythrin from Burkholderia pseudomallei.,* https://doi.org/10.2210/pdb4di0/pdb.

Tickle, I., Flensburg, C., Keller, P., Paciorek, W., Sharff, A., Vonrhein, C. & Bricogne, G. (2018). STARANISO.

Vonrhein, C., Flensburg, C., Keller, P., Sharff, A., Smart, O., Paciorek, W., Womack, T. & Bricogne, G. (2011). Data Processing and Analysis with the autoPROC Toolbox,*Acta Crystallographica Section D* **67**, 293-302.
